# TOGGLe, a flexible framework for easily building complex workflows and performing robust large-scale NGS analyses

**DOI:** 10.1101/245480

**Authors:** Christine Tranchant-Dubreuil, Sébastien Ravel, Cécile Monat, Gautier Sarah, Abdoulaye Diallo, Laura Helou, Alexis Dereeper, Ndomassi Tando, Julie Orjuela-Bouniol, François Sabot

**Affiliations:** DIADE IRD, University of Montpellier, 911 Avenue Agropolis, 34934 Montpellier Cedex 5, France; BGPI CIRAD, INRA, TA A-54/K, Campus International de Baillarguet,34398 Montpellier Cedex 5, France; AGAP CIRAD, INRA, SupAgro, TA A-108/03, 1000 Avenue Agropolis, 34398 Montpellier Cedex 5, France; ADNid-QualTech, 830 Avenue du campus agropolis Baillarguet, 34980 Montferrier-sur-Lez, France; IPME IRD, University of Montpellier, 911 Avenue Agropolis, 34934 Montpellier Cedex 5, France; South Green Bioinformatics Platform, BIOVERSITY,CIRAD,INRA,IRD,SupAgro, Montpellier, France

**Keywords:** Data-intensive analysis, Workow manager, NGS pipeline, parallel computing, high-performance computing, reproducible research

## Abstract

The advent of NGS has intensified the need for robust pipelines to perform high-performance automated analyses. The required softwares depend on the sequencing method used to produce raw data (e.g. Whole genome sequencing, Genotyping By Sequencing, RNASeq) as well as the kind of analyses to carry on (GWAS, population structure, differential expression). These tools have to be generic and scalable, and should meet the biologists needs.

Here, we present the new version of TOGGLe (<u>To</u>olbox for <u>G</u>eneric N<u>G</u>S Ana<u>l</u>ys<u>e</u>s), a simple and highly flexible framework to easily and quickly generate pipelines for large-scale second- and third-generation sequencing analyses, including multi-sample and multi-threading support. TOGGLe is a workflow manager designed to be as effortless as possible to use for biologists, so the focus can remain on the analyses. Pipelines are easily customizable and supported analyses are reproducible and shareable. TOGGLe is designed as a generic, adaptable and fast evolutive solution, and has been tested and used in large-scale projects on various organisms. It is freely available at http://toggle.southgreen.fr/, under the GNU GPLv3/CeCill-C licenses) and can be deployed onto HPC clusters as well as on local machines.

## INTRODUCTION

Advances in Next-Generation Sequencing (NGS) technologies have provided a cost-effective approach to unravel many biological questions, and revolutionized our understanding of Biology. Nowadays, any laboratory can be involved in large-scale sequencing projects, delivering astonishing volumes of sequence data. Although NGS are powerful technologies, they shifted the paradigm from data acquisition to data management, storage and *in fine* biological analyses (1). This intensifies the need for robust and easy-to-use pipelines to perform high-performance automated analyses (2). However, available pipelines depend on the sequencing method used to generate raw data and on the type of analyses to perform (variant calling, GWAS, differential gene expression,…) (3, 4, 5, 6).

To enable processing of large datasets, numerous pipelines frameworks are available, either web-based or command-lines based. The formers, Galaxy (7, 8) or Taverna (9) for instance, can be easily understood by lab biologists as they are available through a graphical web interface. Accessible to non-programmers, they allow adding additional workflows, picking up various pre-packed software, but with limited access to their options. In addition, managing more than a dozen of samples, or having access to different versions of the same software, are not possible without having a high-level access level to the host servers. Thus, they are generally used for small-scale analyses, prototyping or training (10). The second type of frameworks, such as Snakemake (11), Bpipe (12), Ruffus (13) or ClusterFlow (14), targets programmers and expert users. Many of theses tools rely on non-standard, adapted programming languages (e.g. derived either from Python, Bash or Make). Even if bioinformaticians can write complex pipelines using those frameworks, they require high-level skills in programming and are not suitable for lab biologists.

TOGGLe (<u>To</u>olbox for <u>G</u>eneric N<u>G</u>S Ana<u>l</u>ys<u>e</u>s) is an alternate solution mixing advantages from the two kinds of workflow managers, offering a robust and scalable bioinformatic framework for a wide range of NGS applications. Compared to the previous version (version 2 (15)), the current TOGGLe (version 3) is a framework based on command-line, and no more hard-coded pipelines, aimed to both biologists and bioinformaticians. TOGGLe is highly flexible on the data type, working on sequencing raw data (Illumina, PacificBiosciences…), as well as on various other formats (e.g. SAM, BED, VCF) (Figure 1). Carrying out analyses does not require any programming skills, but only basic Linux ones. With TOGGLe, scientists can create robust, customizable, reproducible and complex pipelines, through an user-friendly approach, specifying the software versions as well as parameters.

**Figure 1.**
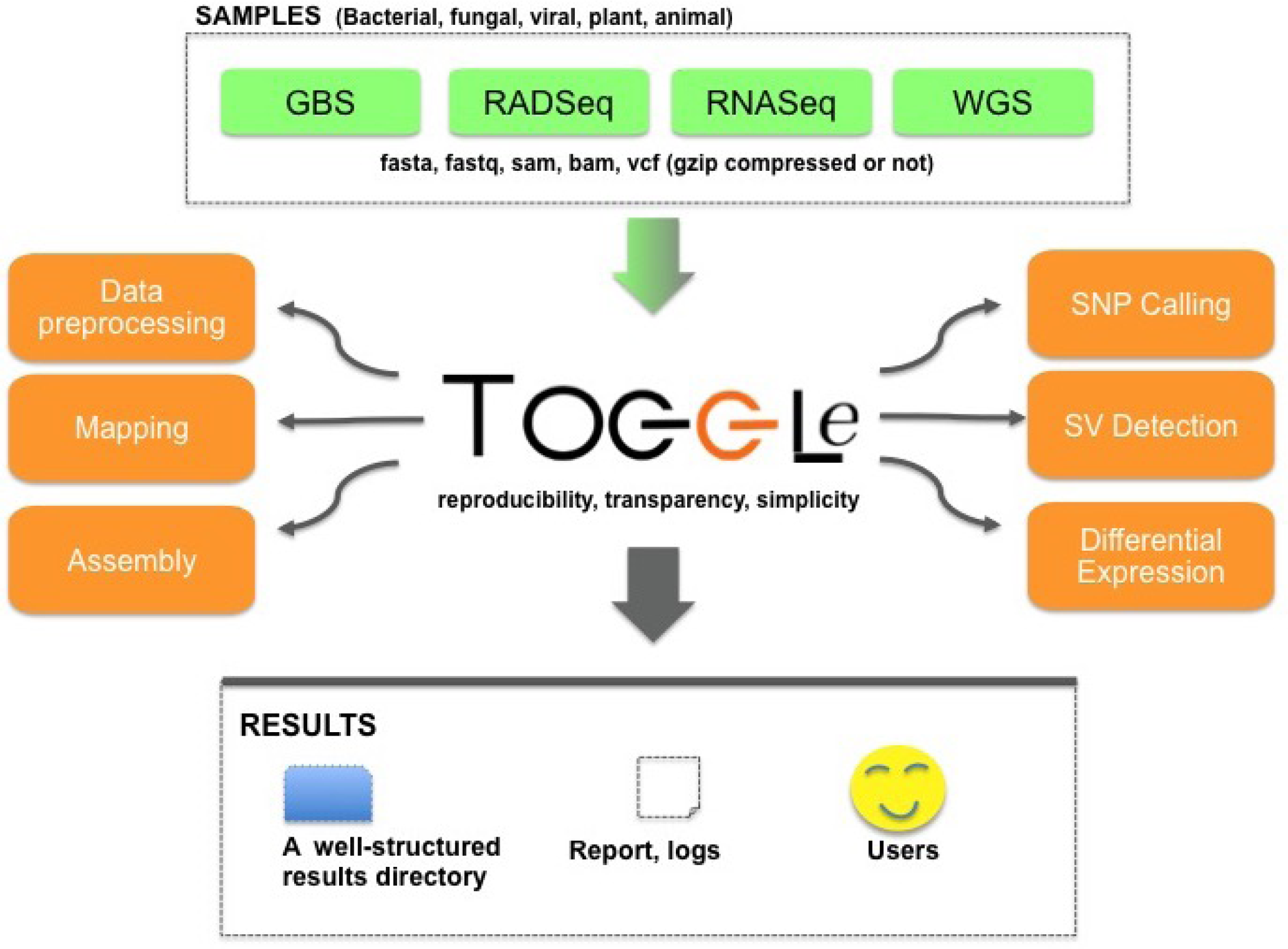
Overview of TOGGLe, a flexible framework for performing large-scale NGS analyses.

## DESIGN AND IMPLEMENTATION

### Input data formats and sample IDs

There is no limit to the number of samples, as soon as all the files are in the same unique directory and of the same format. Input data format can be either FASTA, FASTQ (paired-end, single-end and mate-pair; first-, second- and third-generation), SAM, BAM, BED, GFF or VCF, either plain or compressed (*i.e.* gzip).

Sample IDs and read groups are automatically generated by TOGGLe using the file read name, and no dedicated nomenclature or external sample declaration is needed. For pair-end/mate-pair FASTQ data, no specific name or directory organization is required for pairs to be recognized as such.

### Running a TOGGLe pipeline

TOGGLe workflows can be launch from start-to-end with a single command-line, with three mandatory arguments:

1. the input directory containing files to analyze,
2. the output directory that will contain results generated by TOGGLe,
3. the configuration file.

According to the workflow (e.g. a reads mapping step upon a reference), a transcriptome or genome reference sequence, an annotation file or a key file (for demultiplexing) may be also provided.

### Configuration file

The configuration file plays a central role in TOGGLe, by enabling users to generate highly flexible and customizable workflows in a simple way. Indeed, this basic text file is composed of different parts allowing to build the workflow, to provide software parameters, to compress or remove intermediate data, and to set up a scheduler if needed (see Figure 2 for more details).

**Figure 2.**
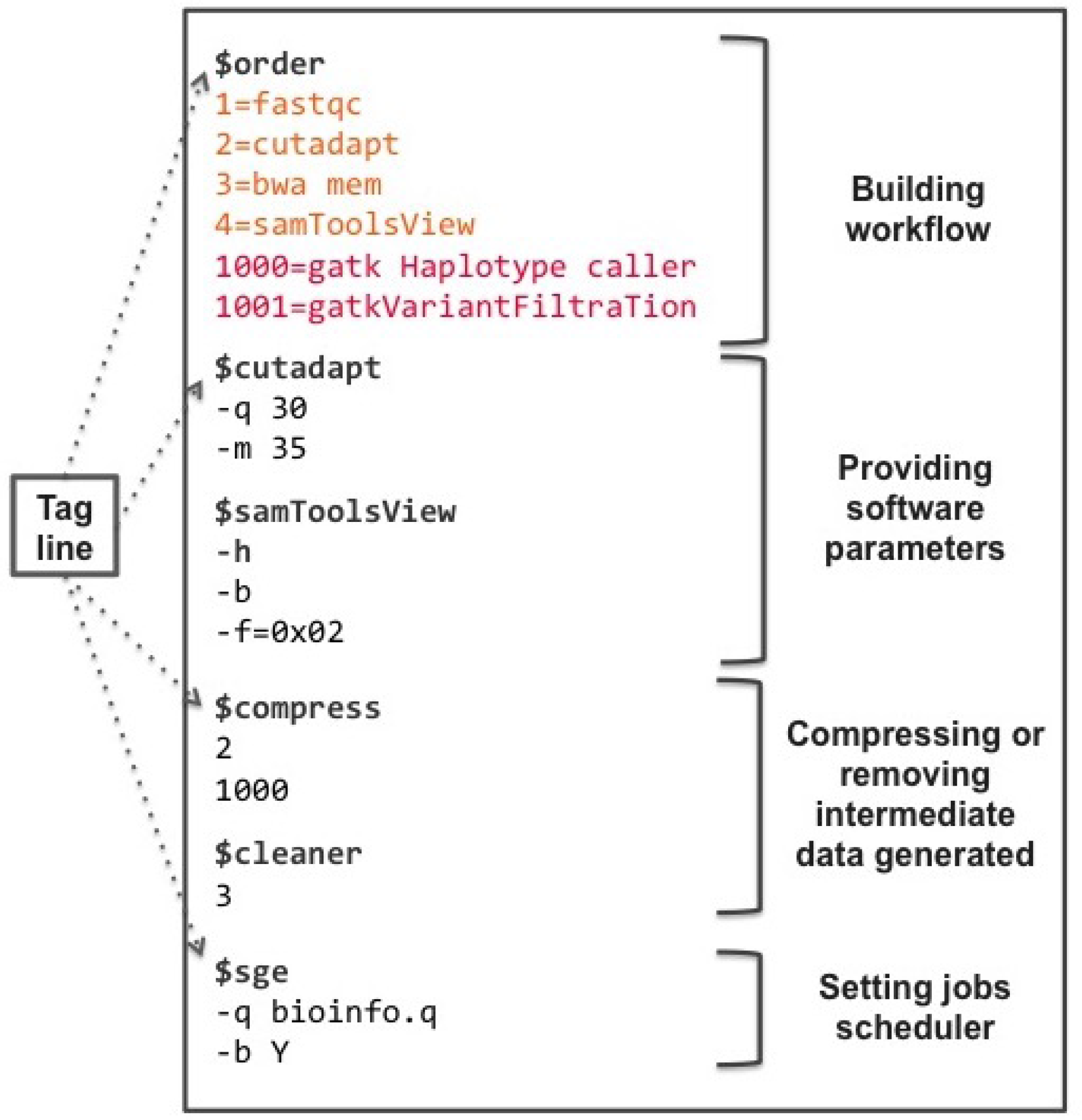
Configuration file to detect polymorphisms, composed of 4 parts : (1) “Building workflow” section : the first 4 analysis (steps lower than 1000, in orange) will be carried out separately for each sample (parallel analyses, in pink) whereas the last two steps will be performed as a global common analysis; (2) “Software parameters”; (3) “Data management”; and (4) “Scheduler management” section

#### Building Workflow

Steps composing the pipeline (e.g. aligning reads upon a reference genome, calling variants, assembling) and their relative order are defined after the *$order* tag. Each line consists of the step number followed by an equal symbol then by the software’s name (e.g. 1=FastQC).

If the step number is lower than 1000, the analysis step is carried out for each sample separately, while the step is performed as a global analysis with the results of all the samples for a value higher or equal to 1000 (see Figure 2 and 3).

**Figure 3.**
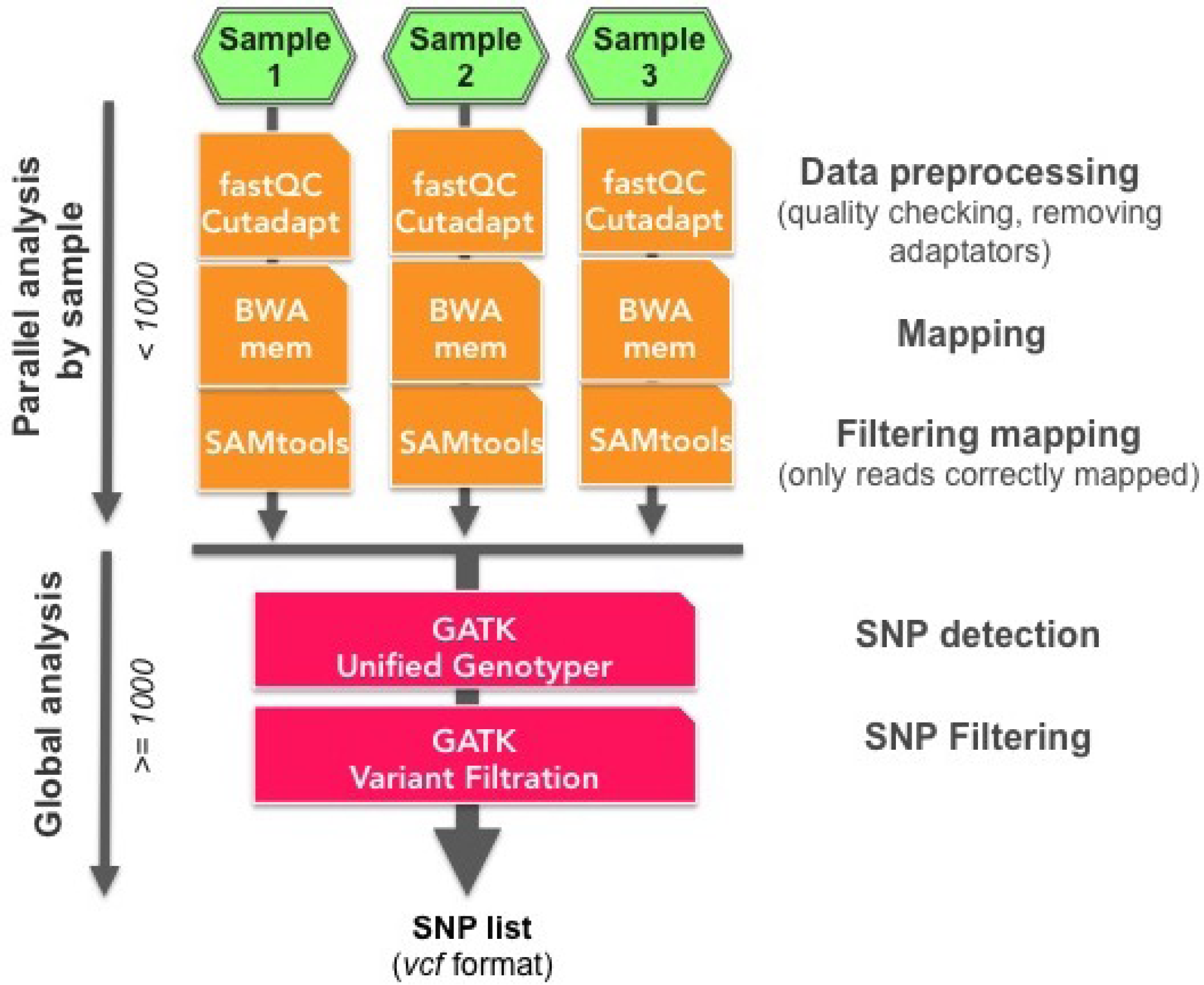
Overview of a TOGGLe pipeline for SNP detection from 3 samples. This figure shows the different steps performed either as parallel (in orange) or as global analysis (in pink). See the corresponding configuration file in Figure 2.

#### Providing software parameters and external tools usage

The syntax for setting software parameters is identical to that used by each software using the command line. These parameters can be provided after the line composed of the symbol *$* followed by the software name (also called tag line; see Figure 2). If no software parameter is provided, the default ones are used. TOGGLe will handle itself the input and output files as well as the references. Users can also use any software not included in TOGGLe with the *generic* tag followed by the command-line.

#### Compressing or removing intermediate data

As analyses generates a large amount of data, we included the possibility to gzip compress or to remove some intermediate data *($compress* and *$cleaner* tags).

#### Setting up jobs scheduler

When analyzing on high performance computing (HPC) systems, TOGGLe runs seamlessly with either LSF, MPRUN, SLURM or SGE jobs schedulers. In addition, users can provide specific environment variables to be transferred to the scheduler (such as the paths or modules to be loaded). Finally, node data transfer is automatically managed by TOGGLe when requested by user.

### Workflows Management

The core of TOGGLe is the *toggleGenerator.pl* script which (*i*) reads the configuration file, (*ii*) generates pipeline scripts, and then (*iii*) executes them as parallel or global analyses (see Figure 4). Basically, *toggleGenerator.pl* acts as a Make-like tool, compiling blocks of code (themselves allowing the execution of the different tools) to create the requested pipeline. It allows the developers to easily add any new tool without having to modify the main code.

**Figure 4.**
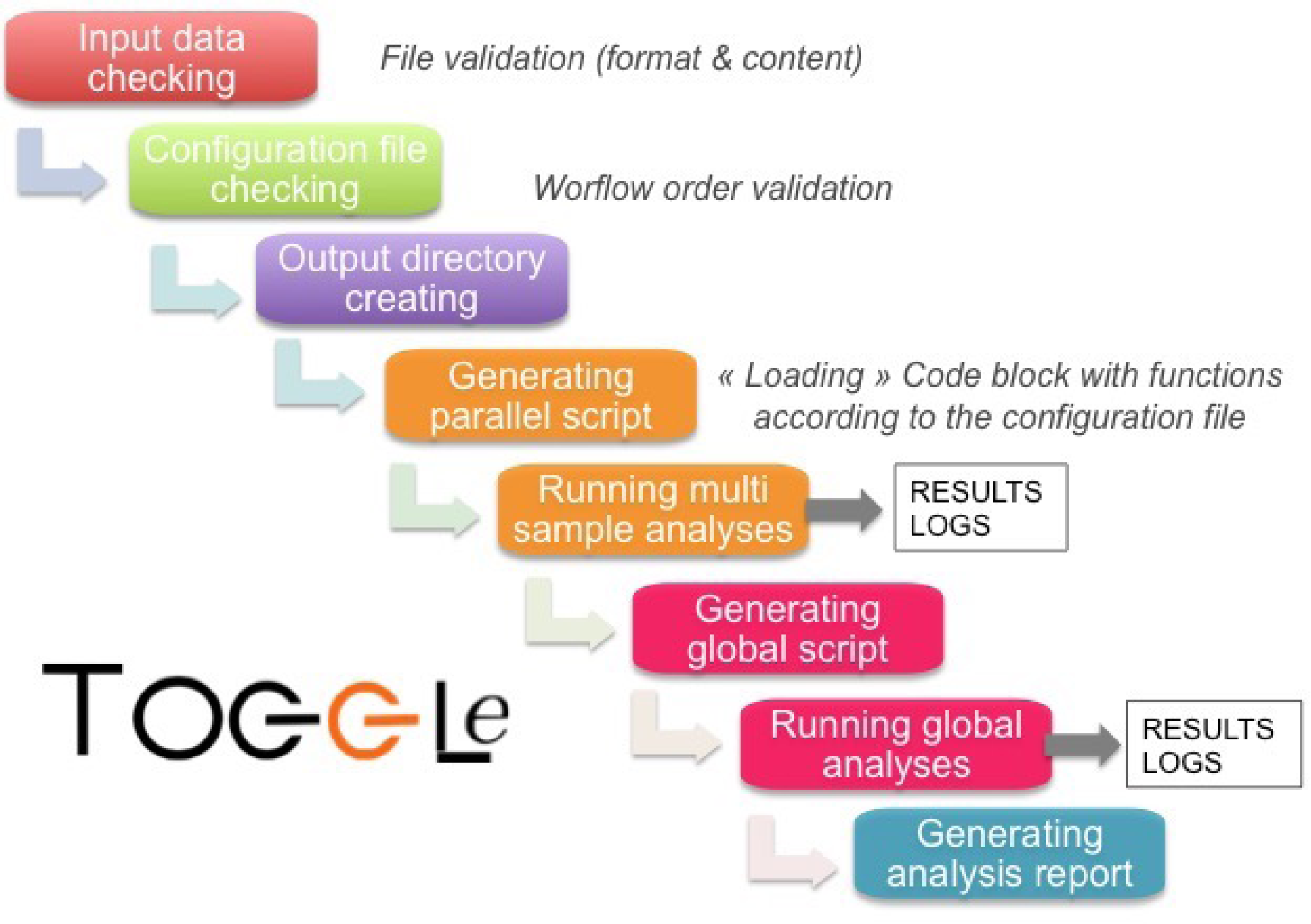
Schematic diagram of the different actions performed by *toggleGenerator.pl.*

### Platforms, Installation and Customization

TOGGLe currently runs on any recent GNU/Linux system (Debian Lenny and more, Ubuntu 12.04 and more, and CentOS 6 and more were tested). TOGGLe was developed to be straightforward to install in several ways : manually (git clone) or through a bash script.

A unique file *(localConfig.pm)* needs to be filled at installation to ensure the integration of the whole software list (path and version; installed separately). However, the whole set of integrated tools is not required to run TOGGLe: one can use it only with SAMtools (16, 17) for instance, and does not need to install the other tools. More detailed information on the different installation procedures can be found at the TOGGLe website (http://toggle.southgreen.fr)

## RESULTS

### Analyses and post-analysis tools integrated in TOGGLe

Developed in Perl, TOGGLe incorporates more than 120 functions from 40 state-of-the-art open-source tools (such as BWA (18), Abyss (19) or FastME (20)) and home-made ones (Table 1 and Supplementary Table S1), making it the command-line workflow manager with the highest number of integrated tools by now (Figure 1; Table 2).

**Table 1.**
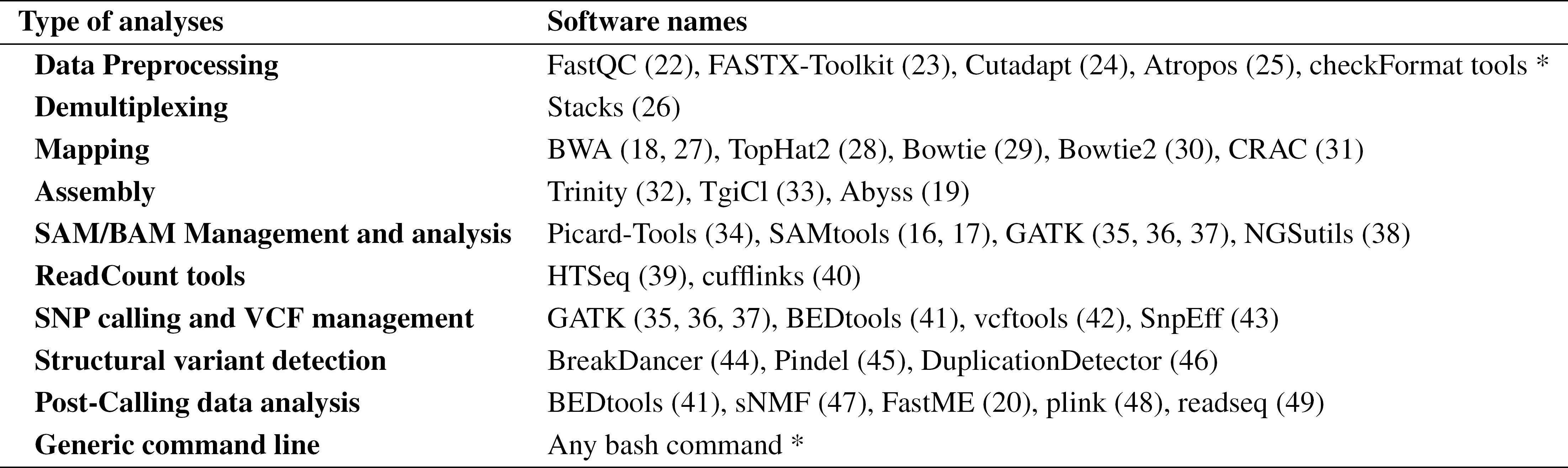
Bioinformatic softwares integrated in TOGGLe. *: home-made tools for data format control and generic command-line.

Once installed, a large array of NGS tools are ready to use with TOGGLe for various type of analyses: input data QC control, cleaning FASTQ, mapping, post-mapping treatment, SNP calling and filtration, structural variation analyses, assembly (genome and transcriptome). A more detailed list of tools implemented is listed in Table 1 and Supplementary Table S1.

Post-analysis tools are available in the current version of TOGGLe, for population genetics, genomic duplication, phylogeny or transcriptomics (Table 1 and Supplementary Table S1).

### A tool targeting both biologists and bioinformaticians

#### Ease of use

TOGGLe drastically simplifies NGS analyses (such as SNP calling, differential expression for RNA, *in silico* assembly) for biologists. Workflows can be easily set up in a few minutes through a unique configuration file, and can be executed through a short command line. In addition, TOGGLe offers access to all parameters without restrictions (or name change) proposed by each software. Finally, users can provide any reference files, without any additional step to add them, at the opposite of ClusterFlow (14) for instance.

As prototyping optimized workflows (software order and parameters) requires a good knowledge of the tools to be used, numerous pre-defined validated configuration files are available on our website (http://toggle.southgreen.fr/) for various type of analyses.

In addition, the effective output files naming convention used by TOGGLe as the well-organized file tree make it easy for the user to identify the different dataset produced (see Supplementary Figure 1).

#### Ease of development and evolution: Simpler is Clever

TOGGLe is designed as a set of separated modules/functions and blocks of code, simplifying code integration and evolution. Each module is written either to run bioinformatic softwares or to ensure functionalities for a specific purpose (such as checking input file formats). The block files are composed of codes implementing a single function at a time. These blocks are then concatenated together following the user pipeline specifications by *toggleGenerator.pl*, to provide a dedicated script pipeline.

This code modularity as well as testing and development processes adopted in TOGGLe prevents the regressions and bugs, facilitating maintenance in a collaborative environment.

#### Production and development versions, and speed of releasing

Two versions of TOGGLe are available, the production one and the development version. The former (http://toggle.southgreen.fr) is validated on a large set of tested data, while the latter (http://github.com/SouthGreenPlatform/TOGGLE-DEV/), more up-to-date, is not completely validated and may provoke errors. Nevertheless, the development version is merged to the production version on a regular basis.

### A robust bioinformatics framework

As TOGGLe was developed initially for performing data-intensive bioinformatics analyses, our main aim was to build a robust workflow framework without sacrificing the simplicity of use and the ease of development (Table 2).

**Table 2.** Comparison with some available command-line workflows managers. Release frequency was estimated from previous release date for each tool. *: no tools are already included in Bpipe and SnakeMake, users have to include them through coding by themselves.

| Features | TOGGLE v3 | TOGGLE v2 (15) | Bpipe (12) | Clusterflow (14) | SnakeMake (11) |
| --- | --- | --- | --- | --- | --- |
| Biologist use | +++ | ++ | - | + | - |
| Ease of pipeline design for biologists | +++ | - | + | ++ | - |
| Transparent supplementary actions | +++ | - | - | - | + |
| Pipeline configuration language | Text | Text | Bash-like | Tabulated format | Python |
| Integrated tools | 120 | 50 | None* | 31 | None* |
| Sanity check | +++ | ++ | + | + | - |
| Software version control | +++ | + | + | - | + |
| Reproducibility | +++ | ++ | ++ | ++ | ++ |
| Re-entrancy | + | - | ++ | + | +++ |
| Schedulers integration | +++ | + | +++ | +++ | ++ |
| Parallel-then-Global analyses | +++ | ++ | - | - | + |
| Easy to install | +++ | ++ | + | ++ | ++ |
| Documentation | +++ | + | ++ | ++ | +++ |
| Release frequency | +++ | - | ++ | + | +++ |

#### Pipeline and data sanity controls

Numerous automatic controls are carried out at different levels as transparent actions : validation of the workflow structure defined by the user (checking if the output file format by step *n* is supported by the step n+1), format control on input data provided by user, checking format of intermediate data.

Missing but requested steps for ensuring the pipeline running (such as indexing reference) are added automatically if omitted.

#### Reproducibility and traceability

TOGGLe ensures that all experimental results are reproducible, documented as well as ready to be retrieved and shared. Indeed, results are organized in a structured tree of directories: all outputs are sorted into separate directories grouped by analyses type (sample or global analyses) and by workflow step (see supplementary figure 1).

All parameters, commands executed as well as software versions are kept in logs and a PDF report. Files such as the pipeline configuration and reference data are duplicated in the output folder, in which are also produced the specific scripts used for the analyses. The original input files, at the opposite to reference and configuration files, are not duplicated to reduce disk usage. Finally, a PDF report for the whole analysis is produced, providing global and visual informations for each sample at each step of the workflow. This report provides a diagram of the workflow, the parameters used (configuration file, softwares version,…) and summary statistics for the key files generated by the pipeline.

#### Error tracking and reentrancy

All errors and warnings encountered are recorded in logs; hence, finding the origin of an error is simplified. If a sample provokes an error, this sample will be ignored for the rest of the analysis, and the failing reported in the error logs. Moreover, as TOGGLe can start from any format or tool, restarting jobs from the failure point is easy and avoid to re-launch the whole analysis.

#### Large numbers of sample analyzed

There is no true limits to the number of samples or the sequencing depth of a project that TOGGLe can take in charge. TOGGLe was already used on hundreds of samples jointly (up to 3,000 by now), from different types of assays (RNAseq, GBS, WGS,…), different analyses (polymorphism detection, read count), and on different organisms (Table 3). The only observed current limits are the data storage and the computing capacities available.

**Table 3.** List of selected projects using TOGGLe.

| NGS type | Organism | Type | Analysis type | Description |
| --- | --- | --- | --- | --- |
| GBS | Date Palm and *Phoenix sp.* | Plant | Polymorphism detection | 242 samples |
| RNASeq | *Arabidopsis thaliana* | Plant | Differential expression | 36 samples, 14 millions reads per sample |
|  | Pacaya | Plant | Transcriptome Assembly | 20 samples, 34 millions reads per sample |
|  |  |  | Polymorphism detection |  |
|  |  |  | Differential expression |  |
|  | *Magnaporthe oryzae* | Fungus | Differential expression | 6 samples, 90 millions reads |
|  | *Burkholderia gladioli* | Bacteria | Differential expression | 6 samples, 90 millions reads |
| WGS | *Coffea canephora* | Plant | Polymorphism detection | 12 samples, 55x mean |
|  | African Rice | Plant | Duplication detection | 16 samples, 35x mean (46) |
|  | African and Asian Rices | Plant | Polymorphism detection | 350 samples, 35x mean |
|  | Rice Yellow Mottle Virus | Virus | Polymorphism detection | 50 samples, 8,000x mean (50) |
|  | Leichmanii | Animal | Polymorphism detection | 15 samples, 20x mean |
|  | *Magnaporthe oryzae* | Fungus | Polymorphism detection | 52 samples, 20x mean |
|  | *Micosphaerella fijensis* | Fungus | Polymorphism detection | 160 samples, 50x mean |
| CaptureSeq | Rice | Plant | Polymorphism detection | 200 targets (genes), 100 samples |
|  | Palm | Plant | Polymorphism detection | 150 targets (800 exons), 120 samples |
|  | *Prunus persica* | Plant | Polymorphism detection | 540 targets (exons), 16 samples |
|  | Human | Animal | Polymorphism detection | 241 targets (8800 exons), 48 samples |

#### HPC and parallel execution

TOGGLe supports different job scheduling systems and can be efficiently run either on a distributed computing system or on a single server (default behaviour).

### Documentation

Installation, quick and complete user manuals, screencasts and a complete developer documentation are available on our website http://toggle.southgreen.fr. In addition, we also provide pre-packed configuration files for different types of classical analyses.

## DISCUSSION & PERSPECTIVES

Nowadays, data processing and analyses are the main bottleneck of scientific projects based on the high-throughput sequencing technologies (1). To deal with this issue, various workflow managers were proposed, with different philosophies. GUI tools (such as Galaxy(7, 8)) provide a web interface to easily and quickly design workflows, being developed for users without any programming knowledge. While very useful for prototyping pipelines with a limited data set and for performing small-scale analyses, they are less suitable to carry out large scale analyses or to create highly flexible pipelines (e.g. access to the whole set of software options). In contrast, many of the recent published workflow managers are aimed to bioinformaticians (e.g. Bpipe (12)), based on an extension of a programming language, allowing to design highly customizable and complex workflows through a command-line interface. However, this efficiency comes at the expense of the ease of use.

TOGGLe is an alternative solution taking advantages of these two kinds of tools. Based on a command-line interface, it is suitable for bioinformaticians as well as biologists. Through a unique text file, it offers a simple way to build highly complex and completely customizable workflows for a broad range of NGS applications, from raw initial data to high-quality post-analyses ones. This ease of use allows to greatly reduce hands-on time spent on technical aspects and to focus on the analyses themselves.

Thus, to improve user experience, TOGGLe performs, in a transparent way, different actions requested to ensure the success of the analysis: automatic reference indexing, data format checking, workflow structure validation, and so on. In addition, the unique possibility offered by TOGGLe to carry out analyses at different levels (per samples then global) reduces the difficulty of use, the manipulation time, and the errors linked to file manipulations (Table 2).

TOGGLe was used on numerous sequencing projects with high number of samples, or high depth sequencing or both (Table 3), computing capacities and data storage being the only observed limits. It was shown to be highly adaptable to various biological questions as well as to a large array of computing architecture and data.

TOGGLe has still some features to be enhanced, as a complete reentrancy after a failed run is not yet possible and request the user to modify the data organization before relaunching. We are thus working on a new management version allowing to re-launch directly without any other manipulations. Moreover, the current version of the configuration file is not exportable to another workflow system using CWL (Common Workflow Language), and it would be interesting to also provide in TOGGLe a translator to CWL. Finally, the current grain of parallelization is the sample level (as for ClusterFlow or Bpipe), and a lot of computation time can also be saved at that point. We are exploring the way to use embarrassingly-parallel approaches (splitting samples in numerous subsamples), in order to run optimized pipelines on highly distributed infrastructures.

As NGS analyses and sequencing technologies are continuously evolving, the modularity and the ease of development of our tool is a true advantage to stay up-to-date. For instance, the two-year old previous version of our software, TOGGLe v2 (15), integrated only 50 tools and was hard-coded. By now, more than 120 tools are available with the current TOGGLe v3, the pipeline creation system is much more flexible and robust, and new tools are regularly under integration to widen the scope of analyses possible using TOGGLe (phylogeny,metagenomics, pangenomics).

## AVAILABILITY

The source code is freely available at http://toggle.southgreen.fr, under the double license GNU GPLv3/CeCiLL-C.

The TOGGLe website comprises a comprehensive step-by-step tutorial to guide the users to install and run the software. An online issue request is also available under GitHub of the South Green platform(21) (http://github.com/SouthGreenPlatform/TOGGLE/issues/) to report bugs and troubleshooting.

- Project home page: http://toggle.southgreen.fr
- Code repository: https://github.com/SouthGreenPlatform/TOGGLE
- Operating system: GNU/Linux (CentOS, Debian, Ubuntu)
- Programming Language: Perl 5.16 of higher (5.24 recommended)
- License: GNU GPLv3/CeCill-C

## ACKNOWLEDGEMENTS

The authors thank users (especially Emira Cherif, Laurence Albar, Remi Tournebize) for testing the first phases of this new version. We also thank Mohamed Kassam for his feedbacks on the TOGGLe usage, and Mathieu Rouard for his useful comments on the manuscript.

## ADDITIONAL FILES

1 – List of tools currently integrated (Table S1)

2 – Example of results file tree created by TOGGLe (Figure S1)

